# Inbreeding depression leads to reduced fitness in declining populations of wild maize

**DOI:** 10.1101/2023.11.20.567972

**Authors:** Aimee J Schulz, David E Hufnagel, Paul Gepts, Matthew B Hufford

## Abstract

Crop wild relatives can serve as a source of variation for the genetic improvement of modern varieties. However, the realization of this genetic potential depends critically on the conservation of wild populations. In this study, five populations of *Zea mays* ssp. *parviglumi*s, the closest relative of domesticated maize, were collected in Jalisco, Mexico and planted in a common garden. Eleven traits related to plant fitness were measured and evaluated in the context of genetic diversity and genetic load. Plants whose seed were sourced from larger, less disturbed populations had greater genetic diversity, lower genetic load, and possessed phenotypes associated with higher fitness, while plants sourced from smaller, heavily impacted populations had traits characteristic of lower fitness and increased genetic load. For example, plants from larger populations germinated more quickly, reached anthesis sooner, demonstrated a higher level of photosynthetic activity, and produced more above-ground biomass, suggesting a direct correlation between the fitness of a population, genetic diversity, and genetic load. These results emphasize the importance of preserving the habitat of populations of *Zea mays* ssp. *parviglumis* to limit inbreeding depression and maintain the genetic diversity and adaptive potential of this germplasm.

## Introduction

Crop domestication occurred in multiple centers of origin across the world approximately 10,000 years ago (Piperno et al., 2009; Matsuoka et al., 2002; van Heerwaarden, 2011; Larson et al., 2014; Fuller et al., 2014). Although these events took place independently, ancient agriculturalists selected similar traits to meet their needs for food, fuel, and fiber. Traits selected across multiple domesticates include reduced tillering and branching, loss of seed shattering, increased size of harvested plant parts, and reduced seed dormancy (Gepts, 2014). Collectively, these traits comprise what is known as the “domestication syndrome” (Hammer, 1984) and can be detrimental in the wild due to the increased dependence they cause on humans for survival (Darwin, 1859).

Demographic bottlenecks and targeted selection during domestication resulted in a reduction of genetic diversity in crops when compared to crop wild relatives (CWRs). In maize, Hufford *et al*. (2012) found a 20% reduction in nucleotide diversity after domestication. Similar trends have been observed in common bean and soybean with a ∼10-72% and 36% reduction of genetic diversity in each crop respectively versus their wild progenitors (Kwak and Gepts, 2009; Bitocchi, 2013; Lam, 2010). At least a portion of the diversity retained in CWRs and lost in crops could provide useful adaptations for crop improvement (Bohra, 2022; Kapazoglou, 2023; Renzi, 2022). Already, CWRs have meaningfully contributed to breeding programs (Kashyap, 2022; Tirnaz, 2022). For example, in wheat, a salt-tolerance allele from *Triticum monococcum* was successfully introgressed into commercial durum wheat, resulting in a 25% increase in yield when grown under saline conditions (Munns, 2012). In common bean, wild *P. vulgaris* contributed seed weevil resistance (Kornegay *et al*., 1993) attributable to the APA (arcelin, phytohemagglutinin and α-amylase inhibitor) seed protein (Osborn *et al*., 1988; Kami *et al*., 2006). In potatoes, multiple wild diploid *Solanum* species have been used to increase cultivated potato resistance to cold sweetening (Hamernik, 2009). Blight resistance in maize was developed from a resistance allele found in *Tripsacum dactyloides L.* (Goodman, 1987). Additionally, Flint-Garcia *et al*. have found evidence suggesting alleles from the wild relative *Zea mays* ssp. *parviglumis* (hereafter *parviglumis*) could increase protein content in maize kernels (Flint-Garcia, 2009).

While the potential benefits of conserving CWRs are clear, they are vastly underrepresented in international germplasm banks (Castañeda-Álvarez et al., 2016), comprising only 2-18% of total collections (FAO, 2017). Moreover, few conservation efforts and *in situ* protection measures have been put into place to specifically protect CWRs (Wilkes, 2007; Castañeda, 2016; Goettsch, 2021). This is especially true for teosinte, the common name for maize wild species in the genus *Zea*. The perennial teosintes *Z. perennis* and *Z. diploperennis* are the only formally protected species, with a study by Sánchez González and co-authors showing that only 11.2% of teosintes are located in Protected Natural Areas (Sánchez González, 2018). In some areas, populations of *parviglumis*, the wild progenitor of maize, may be in serious decline (Wilkes, 2004; Wilkes, 2007). Recent anthropogenic impacts and environmental pressures in southwestern Mexico have resulted in a loss of *parviglumis* habitat and a reduction in gene diversity (Hufford, 2010). Many abandoned maize fields have been converted to rangeland and cash crop fields (Nadal, 2002). Additionally, recent modernization of agriculture, rural livelihood diversification, rural-urban and translational migration, and livestock intensification threaten traditional agricultural systems and maize landraces (Keleman, 2009). Traditional agricultural systems may be integral for teosinte, as *parviglumis* is a ruderal species and dependent on the intermediate levels of habitat disturbance these systems provide (Sanchez-Velasquez, 2002).

In this study, five *parviglumis* populations were sampled from the western part of the state of Jalisco, Mexico, genotyped, and grown in a common garden to evaluate their fitness. The five populations were named for their nearest town: Ejutla A (EJUA), Ejutla B (EJUB), La Mesa (MSA), San Lorenzo (SLO), and El Grullo (ELG). These populations show a broad range of genetic diversity and we suspected that smaller, more fragmented populations may be experiencing inbreeding depression. We hypothesized that populations with the least genetic diversity would also have the lowest fitness (*i.e.*, lower plant height, fewer tillers, less above-ground biomass, longer days until flowering) and the highest levels of genetic load. If verified, our findings will help characterize the conservation status of *parviglumis* populations in Jalisco, Mexico, and allow for a better assessment of conservation needs.

## Methods

### Sampling of *parviglumis* populations

The samples grown in the common garden were collected from five *parviglumis* populations in the southwestern state of Jalisco, Mexico (Table 1). These populations were found 20km east of Autlán de Navarro in an area spanning 25km east to west between the southeastern town of El Grullo and northeastern town of San Lorenzo. The first population was a small roadside population of less than 100 *parviglumis* plants near the town of El Grullo (ELG). The EJUA population consisted of several million, high-density *parviglumis* plants. This population was the most genetically diverse of those sampled and had minimal observed anthropogenic impacts. The third population, Ejutla B (EJUB), was located 5km from EJUA on the opposite side of the village. Ejutla B occurred in a mixed-use landscape, consisting of lightly forested areas, rangeland, and maize cultivation. Farmer interviews from this area revealed a recent reduction in population size and introduction of exotic forage grasses. The fourth population was found on a plateau near the town of La Mesa (MSA). This population was widely dispersed and found both in the margins of row-cropped maize and along roadsides. Finally, the fifth population was found near the town of San Lorenzo (SLO) among traditional subsistence-level maize cultivation. Due to the widespread planting of non-native forage grasses (*Brachiaria* sp.), the population size had been greatly reduced.

**Table 1:**
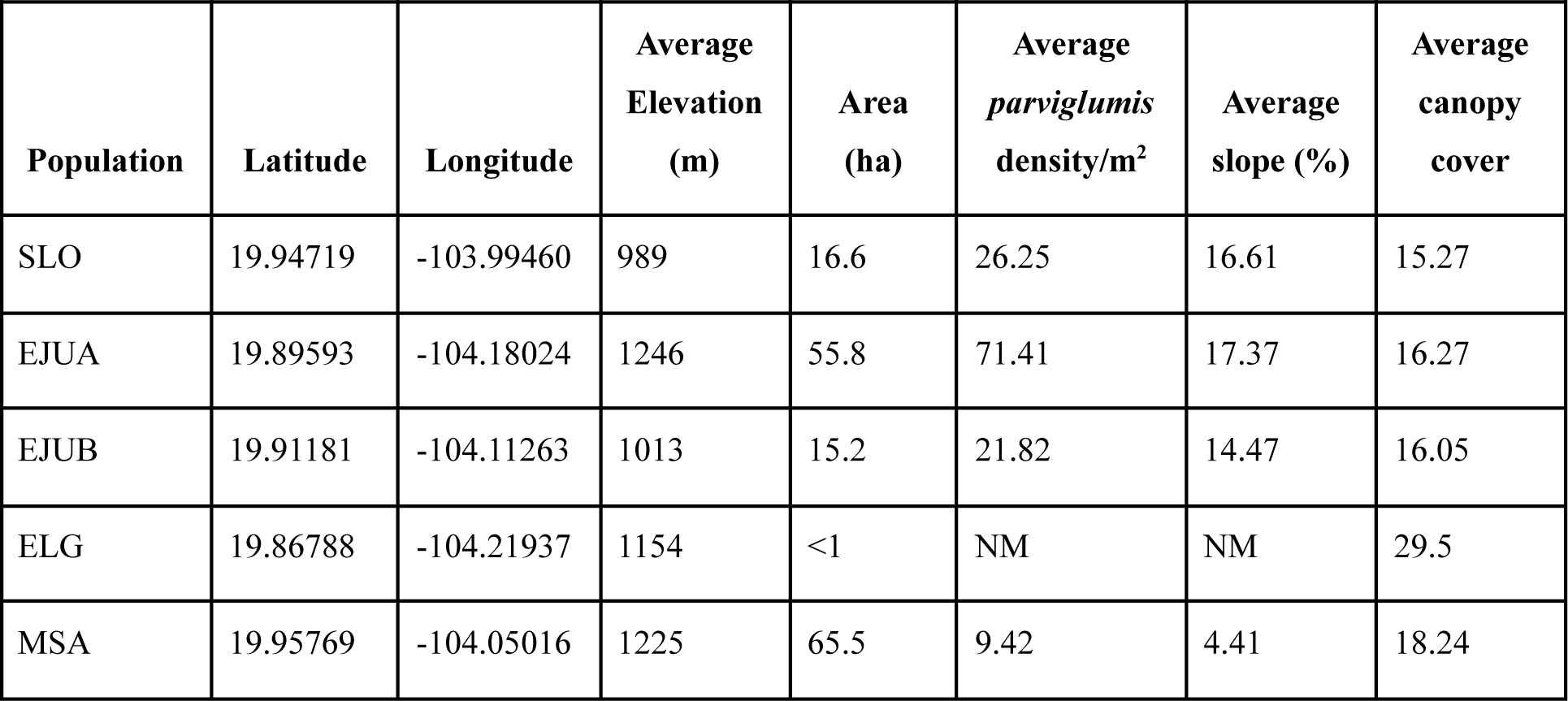
Information on each of the five *parviglumis* populations collected.

Each population was mapped with a GPS receiver during February, March, and September of 2007. Based on the population maps, virtual transects were created and used to collect a random, stratified sample of the five populations. Collections were made at approximately 50 sampling sites per population. Due to the small population size, ELG had only 26 sampling sites. At each sampling site, seed from 10 individuals within a 10-m radius were collected and pooled. A P587 permit from USDA-APHIS was obtained for planting and propagation of seeds. The seeds were then transported to the field site in Davis, California, for a common garden experiment.

### Common Garden Design

The common garden was located at the University of California Plant Sciences Field Facility in Davis (38.532662°N. Lat.; 121.769954°W. Long.). Due to *parviglumis* being day length sensitive, the samples were planted in mid-July so that they would be at an appropriate growth stage to flower in early October when northern California reaches short days. Five complete blocks were planted with three seeds from each of the 50 sampling sites of the four larger populations. Twenty-six seeds from the ELG population were planted in each block due to sample size limitations. Planting locations for the seeds were randomized in each of the blocks. Prior to planting, the seeds were scarified with 20% hydrogen peroxide for five hours. Seeds were spaced at 30cm during planting. To minimize edge effects, a two-row buffer of a commercial maize F_1_ hybrid surrounded the entire garden. Fertilizer was also incorporated into the field before planting. Seed weight (SW) was measured before planting to account for maternal effects. Days to Germination (DTG) was measured during the first two weeks after planting and again one month after planting. Plant height was measured at 15 days (PH15), 30 days (PH30), and 60 days (PH60) after germination. Two months after planting, the number of tillers (NT) per plant was recorded. Once plants reached flowering, the rate of stomatal conductance (CON) was measured using a Decagon Devices leaf porometer, with temperature also measured as a covariate. The leaf porometer measures transpiration over a 30-second period and is an indirect measure of photosynthesis. However, it has been shown in wheat that adaptive traits such as water use efficiency can also result in decreased transpiration, so CON should be interpreted carefully (Yuping, 2017). The number of days until the start of flowering (first emergence of a tassel, DTF) was recorded for each plant. A subsample of 10 individuals per population per block was used for determining the pollen viability (PV) of the population. A fluorescein diacetate assay (Heslop-Harrison 1970) was used to determine the proportion of stainable pollen grains out of 200 pollen grains per plant. Plant survival (PS) was measured for each population upon the conclusion of the experiment. Plants were then placed for two weeks in a drying oven, and subsequently, measured for total aboveground biomass (TAB). An onset of frost before seed maturation resulted in the seed set not being determined.

### DNA extraction, polymerase chain reaction, and genotyping

Dried leaf samples from 201 teosinte plants were ground using a Mini-Beadbeater-96 homogenizer (Biospec Products) and 3mm borosilicate beads. The genomic DNA from the ground samples was extracted using a modified CTAB protocol based on Saghai-Maroof et al. (1984) and concentration measured using a Hoechst 33258 Assay on a DyNA Quant 200 fluorometer. The genomic DNA was then diluted to a 10 ng/μl working solution for polymerase chain reaction (PCR). A total of 18 SSRs were selected from the 10 linkage groups to ensure adequate coverage (Supplemental Table 1), and neutral markers were selected from SSRs developed for maize. The neutral markers were unlinked, consisted of tri-nucleotide repeats, and a PIC > 0.5.

A PCR protocol based on Schuelke 2000 was implemented, where a M13 reverse sequence tail (TGTAAAACGACGGCCAGTATGC) was added to the 5’ end of the forward primers and fluorescent labels, resulting in no need to directly label each forward primer. Each PCR reaction was 20μl, consisting of 30ng DNA, 32 pmol M13-labeled 6-FAM fluorescent dye (Sigma Life Science), 8 pmol M13-labeled forward primer and 32 pmol reverse primer (Sigma Life Science), standard Taq buffer (New England Biolabs), 0.5 unit of Taq polymerase (New England Biolabs), and 0.2 μM dNTP (New England Biolabs). The PCR cycles were as follows: five minutes at 94°C, 30 cycles of 30 seconds at 94°C, 45 seconds at 56°C, and 45 seconds at °C, 8 cycles of 30 seconds at 94°C, 45 seconds at 53°C, and 45 seconds at 72°C, and a 10-minute extension at 72°C. The amplified fragments were diluted to 1/10th or 1/20th of their initial concentration using sterile water and were multiplexed according to their size. 2 μl of each diluted and multiplexed PCR product was then added to 10 μl of Hi-Di Formamide and 0.1 μl of GeneScan 500 LIZ Size Standard (Applied Biosystems Inc.) to prepare for genotyping. These were then denatured for 3 minutes at 95°C and analyzed on an ABI 3730 instrument (Applied Biosystems Inc.) at the Veterinary Genetics Laboratory at the University of California, Davis. GeneMarker (version 1.85, SoftGenetics LLC, State College, PA) was used to score genotypes.

### Genetic Diversity

Input files were prepared for downstream analysis using CREATE 1.33 (Coombs et al., 2008). Using GENALEX 6 (Peakall and Smouse, 2006), genetic diversity within each population was estimated for the expected (HE) and observed (HO) heterozygosities, number of alleles per locus (A), and number of private alleles (AP). To account for sample size differences among populations, allelic richness (AR), or the average number of alleles per locus, was estimated using rarefaction (Hurlbert, 1971) in the software package, FSTAT 2.9.3 (Goudet, 1995). The means across loci for genetic diversity stats were compared using JMP 8 (SAS Institute, Inc.) with Tukey-Kramer HSD test (α = 0.05)and a one-way ANOVA. ARLEQUIN 3.11 (Excoffier et al., 2005) was used to test for departure from HWE for each locus in each population with a test similar to Fisher’s exact test using a Markov chain of 1,000,000 iterations and 100,000 dememorization steps. To correct for multiple tests, a sequential Bonferroni correction at α = 0.05 was used (Rice, 1989). Due to being monomorphic for the majority of loci, the ELG population was not tested for HWE departure like the rest of the populations.

### Statistical Analysis

All statistical analyses were completed in R version 3.4.4 (R Core Team, 2022). Population means were compared for each trait using a mixed-effects ANOVA where data had a normal distribution or could be transformed as determined by a Kolmogorof-Smirnoff test. Significant differences between the populations for trait means were determined using a Tukey HSD test. The four larger populations (EJUA, EJUB, MSA, SLO) were first analyzed using multiple sampling sites as a randomized complete block design with five replications in a mixed effects ANOVA. Block and sampling site were random effects, and population was a fixed effect. To compare the four larger populations with the smaller ELG, trait values were averaged across all sampling sites for each population and then analyzed as a randomized complete block design without replication. Block was considered a random effect, and population a fixed effect in the ANOVA. Additionally, a Levene’s test (Fox, 2019) for homogeneity of variance was conducted for each analysis. Gene diversity and allelic richness were evaluated using microsatellite data with RStudio version 3.4.4 (Rstudio Team, 2020) in the packages adegenet (Jombart, 2008; Jombart, 2011), pegas (Paradis, 2010), and hierfstat (Goudet, 2005). Principal component analysis of the microsatellite data was completed using the R package ade4 (Chessel, 2004). When determining the correlation between genetic diversity and population fitness, the trait means were scaled so that 1 was the fittest population and 0 the least fit population.

### Population Structure

Microsatellite data of the five populations were analyzed using STRUCTURE (Pritchard, 2000; Pritchard, 2007) consisting of 17 loci and 201 individuals. A burnin of 500,000 followed by a run length of 300,000 was used on an ancestry model with an initial alpha of 0.3333. An individual alpha was generated for each population due to the unequal sample size. The best k value was estimated using posterior probability through Clumpak (Itay, 2015). StructureHarvester (Vonholdt, 2012) was also utilized for estimating k values.

### Genetic Load

Single nucleotide polymorphism (SNP) data (Pyhäjärvi *et al*. 2013) from 48 *parviglumis* individuals from four of our populations were used for genetic load analysis. The format was converted from TopStrand to Ref/Alt using a custom Python script, and loci names were converted from AGPv2 to AGPv3. Alleles at each SNP were then compared to the respective derived allele and adjusted GERP score for that locus (Wang, 2017; Rodgers-Melnick, 2015). Genetic load under the recessive model was determined by summing the GERP scores for all homozygous derived alleles for each individual. Load under the additive model was calculated by summing the GERP score of every derived allele (heterozygous or homozygous) for each individual. The resulting data were evaluated for normal distribution prior to completing an ANOVA. Homogeneity of variance was checked through a Levene’s test. A Tukey HSD test was used to compare between the populations.

## Results

### Population Structure

Our five populations were distributed across 25km and are geographically distinct (Figure 1A), indicating gene flow may be limited between populations and providing the opportunity for genetic structure to accumulate. To gauge this, a principal component analysis of the microsatellite data showed an initial clustering of the five populations into three distinct genetic groups: SLO and MSA, EJUA and EJUB, and ELG (Figure 1B).

**Figure 1:**
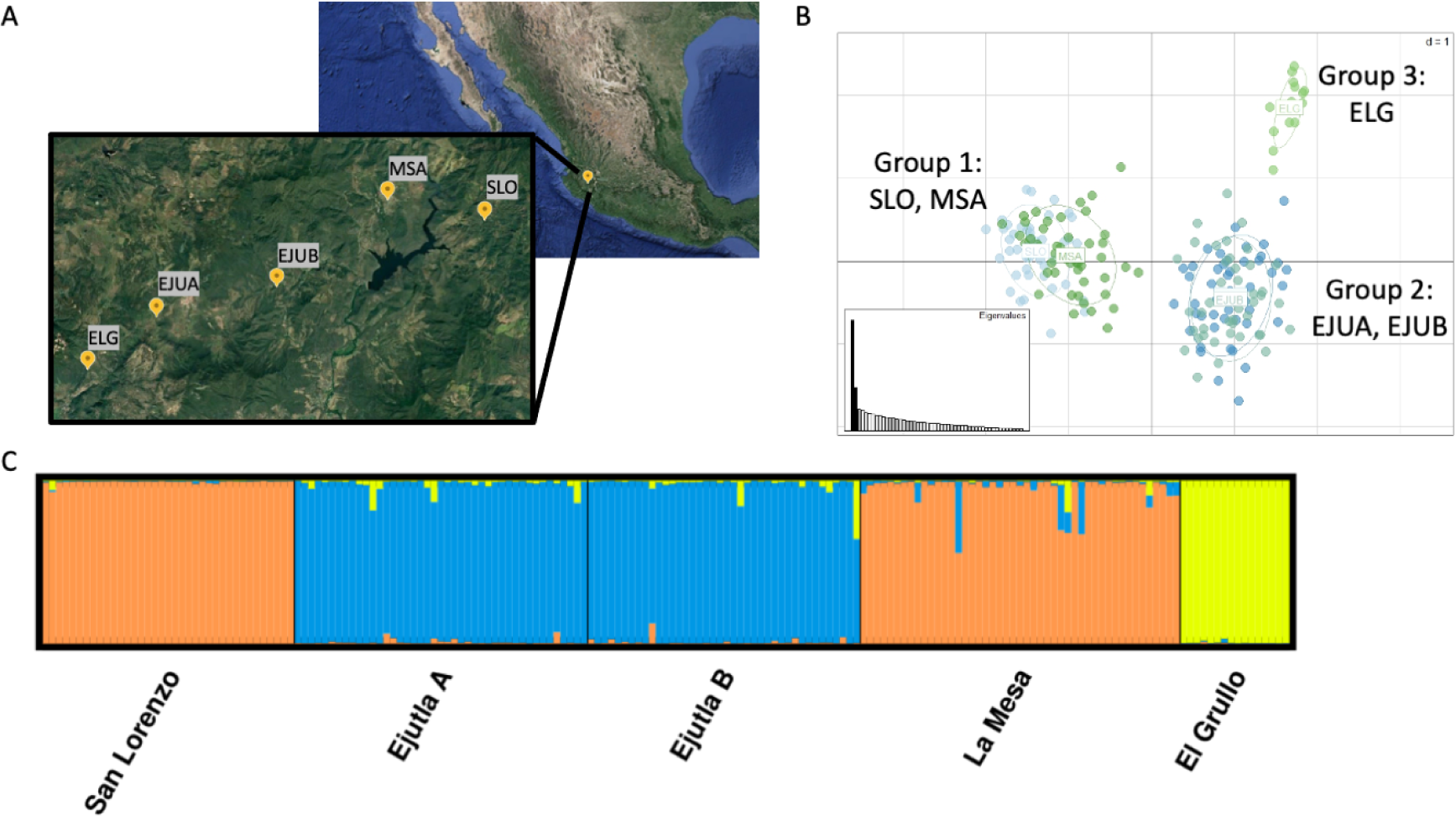
A) Location of the five teosinte populations. B) PCA of the five populations using microsatellite data. C) STRUCTURE plot of the five populations, k = 3.

Structure was used to further confirm the population structures of the five groups (Figure 1C), where k = 3 was determined to be the most appropriate. EJUA and EJUB formed one population, while SLO and MSA grouped into the other. ELG was genetically distinct from the other two population groups under a k = 3 model but grouped with EJUA and EJUB in the k = 2 model.

### Population Genetic Diversity

Microsatellite data from 201 individuals and 17 loci were analyzed to determine the degree of genetic diversity in each population. EJUA and EJUB had the highest expected heterozygosity, with MSA at an intermediate level and SLO and ELG at the lowest levels. When comparing observed heterozygosity, EJUA had the highest levels, while EJUB and MSA both had intermediate levels. SLO and ELG had the lowest levels of observed heterozygosity. The five populations followed similar trends with allelic richness (Table 2).

**Table 2:**
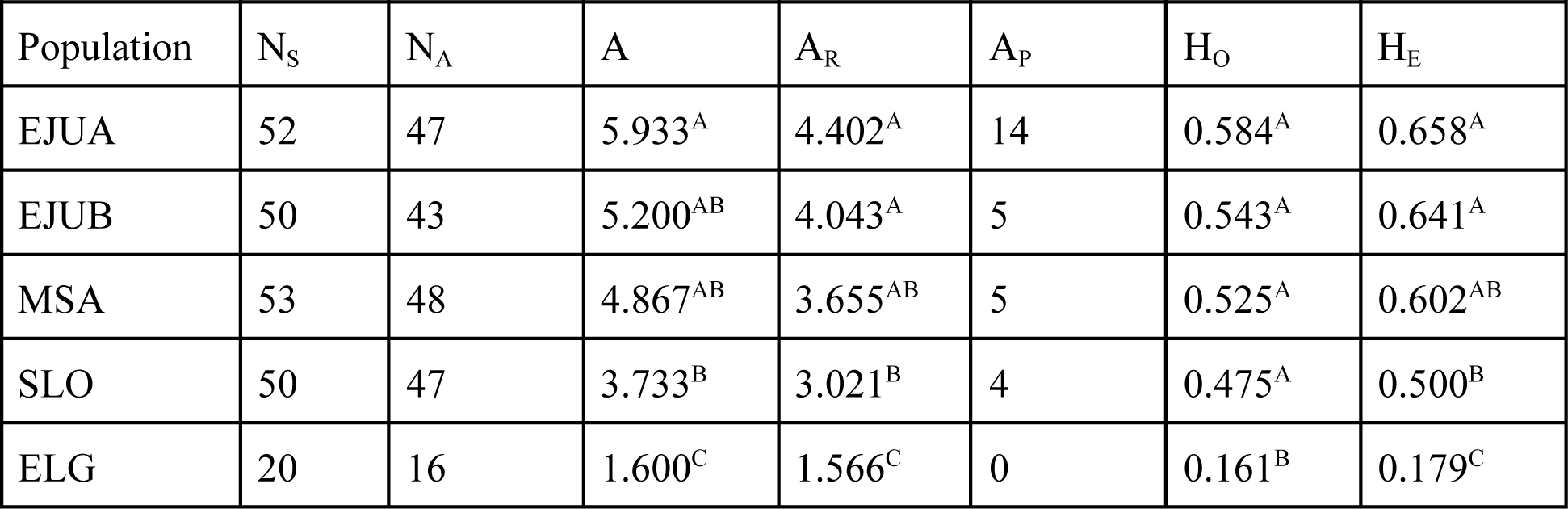
Genetic diversity of the five populations using 15 neutral microsatellite loci. Superscripted letters indicate statistically significant groups based on a Tukey-Kramer HSD test. N_S_, number of samples collected; N_A_, number of samples included in analysis; A, average number of alleles per locus; A_R_, average allelic richness; A_P_, number of private alleles; H_O_, average observed heterozygosity; H_E_, average gene diversity

| Population | N<sub>S</sub> | N<sub>A</sub> | A | A<sub>R</sub> | A<sub>P</sub> | H<sub>O</sub> | H<sub>E</sub> |
| --- | --- | --- | --- | --- | --- | --- | --- |
| EJUA | 52 | 47 | 5.933<sup>A</sup> | 4.402<sup>A</sup> | 14 | 0.584<sup>A</sup> | 0.658<sup>A</sup> |
| EJUB | 50 | 43 | 5.200<sup>AB</sup> | 4.043<sup>A</sup> | 5 | 0.543<sup>A</sup> | 0.641<sup>A</sup> |
| MSA | 53 | 48 | 4.867<sup>AB</sup> | 3.655<sup>AB</sup> | 5 | 0.525<sup>A</sup> | 0.602<sup>AB</sup> |
| SLO | 50 | 47 | 3.733<sup>B</sup> | 3.021<sup>B</sup> | 4 | 0.475<sup>A</sup> | 0.500<sup>B</sup> |
| ELG | 20 | 16 | 1.600<sup>C</sup> | 1.566<sup>C</sup> | 0 | 0.161<sup>B</sup> | 0.179<sup>C</sup> |

### Germination

All populations had relatively high germination rates. EJUA, the most genetically diverse population (Table 2) had 91 percent germination. EJUB, ELG, and MSA had intermediate germination rates between 84 and 89 percent. SLO had the lowest germination at 82 percent. Significant differences (p = 0.01078) were seen between all populations in the number of days until germination. EJUA germinated markedly sooner than the other populations, with an average time of 6.7 days. The less diverse SLO was the latest to germinate, averaging around 8.5 days.

### Vegetative Traits

#### Plant Height

Among the vegetative traits recorded, significant differences were seen in plant height as early as 15 days after germination (p = 2.198^-07^). These trends continued for 30 days (p = 2.367^-08^) and 60 days (p = 6.832^-07^) after germination. The EJUA and MSA populations were consistently the tallest, with the less diverse SLO and ELG populations considerably shorter throughout the field experiment. EJUB had an intermediate height despite having the second-highest level of genetic diversity.

#### Number of Tillers

The number of tillers, or basal branches, was counted on each plant. EJUA produced significantly more (p = 7.549^-07^) tillers than the SLO and ELG populations. EJUA averaged 9.6 tillers per plant. MSA and EJUB produced 8.2 and 8.4 tillers per plant, respectively.

SLO and ELG produced the least, with the trait means of 7.1 and 5.7 tillers per plant, suggesting plants from these populations were less vigorous.

#### Total Aboveground Biomass

The total aboveground biomass displayed similar trends to other vegetative traits (Figure 2A). EJUA, EJUB, and MSA produced the highest, as would be expected, given their high level of genetic diversity. Significant differences in biomass production were seen between all populations (p = 1.426^-07^). Perhaps most striking, SLO individuals produced half the biomass of EJUA, and the ELG population produced only one-third of the biomass of the EJUA population.

**Figure 2:**
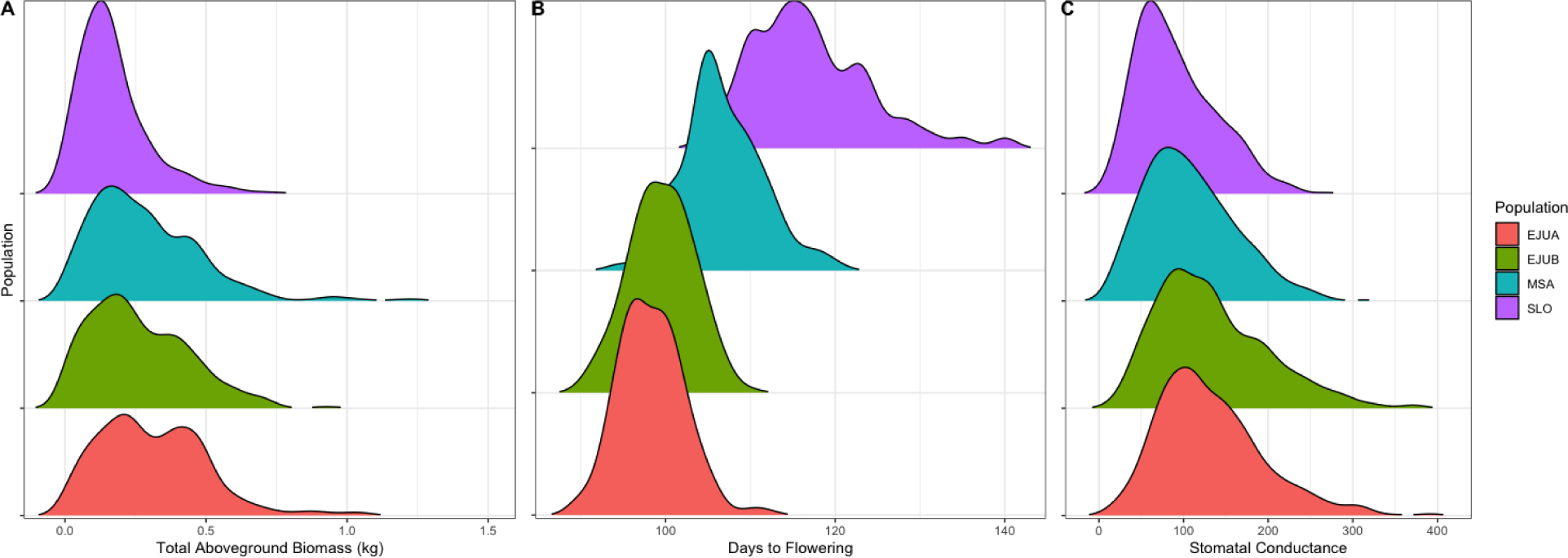
Distribution of the following traits in the four larger populations (EJUA, EJUB, MSA, and SLO): A) total aboveground biomass, B) days to flowering, C) stomatal conductanc.

### Reproductive Traits

#### Days to Flowering

Interestingly, the trends for days to flowering appeared to be separated by geographical region, which also followed the population structure groupings (Figure 2B). The plants in the western region of the sampling area (EJUA, EJUB, and ELG) flowered earlier than the plants sampled from the eastern region (MSA and SLO). Significant differences (p = 1.257^-13^) were seen between all populations for flowering time, and EJUA flowered on average 20 days sooner than SLO. Additionally, SLO had the widest range of flowering time.

#### Pollen Viability

Pollen viability was determined using a fluorescein diacetate assay and measured as percent viable pollen per 200 grains. The percentage of viable pollen was low across all populations, ranging from 0.11-0.22%. Many plants flowered in late November and early December, and we suspect that the cold temperatures of Northern California were causing low pollen viability. There were significant differences between the populations for pollen viability, but regional trends similar to days to flowering were seen between EJUA (0.22% viable) and SLO (0.11% viable).

#### Stomatal Conductance

Stomatal conductance showed similar trends as the number of days to flowering (Figure 2C). Once again, the western region (EJUA, EJUB, and ELG) appeared to have higher overall conductance than the eastern region (MSA and SLO). As with days to flowering, the populations with lower genetic diversity had the lowest stomatal conductance. There was a significant difference between all populations in stomatal conductance (p = 1.902^-08^). Since stomatal conductance is linked to photosynthetic rate, populations with lower stomatal conductance can be inferred to have an overall lower photosynthetic rate.

#### Survival

The survival rate across all populations was quite high, ranging from 95 to 99 percent. The EJUA and EJUB populations had the highest survival rates at 99%. The MSA and SLO populations had intermediate survival rates at 97%. Finally, the ELG population had the lowest survival rate, with 95% of plants making it through the end of the experiment.

### Scaled Fitness and Gene Diversity

All trait means were scaled for each population with 1 being the fittest and 0 being the least fit. When compared with the estimated expected heterozygosity (He; gene diversity, Table 2) in the same populations, we saw a consistently positive correlation between gene diversity and fitness (Figure 3). The vegetative traits (PH15, PH30, PH60, TAB, and NT) saw a more direct correlation, with R^2^ ranging from 0.61 to 0.89 (Supplemental Figure 1). Days to flowering, pollen viability, and stomatal conductance were less strongly correlated. This was due to the ELG population having a higher fitness than expected in these traits despite a lower overall gene diversity. However, when ELG was removed and the trait means from the four larger populations were compared, the correlation between gene diversity and fitness became much clearer (i.e., the R^2^ for DTF increased from 0.07 to 0.99) (Supplemental Figure 2).

**Figure 3:**
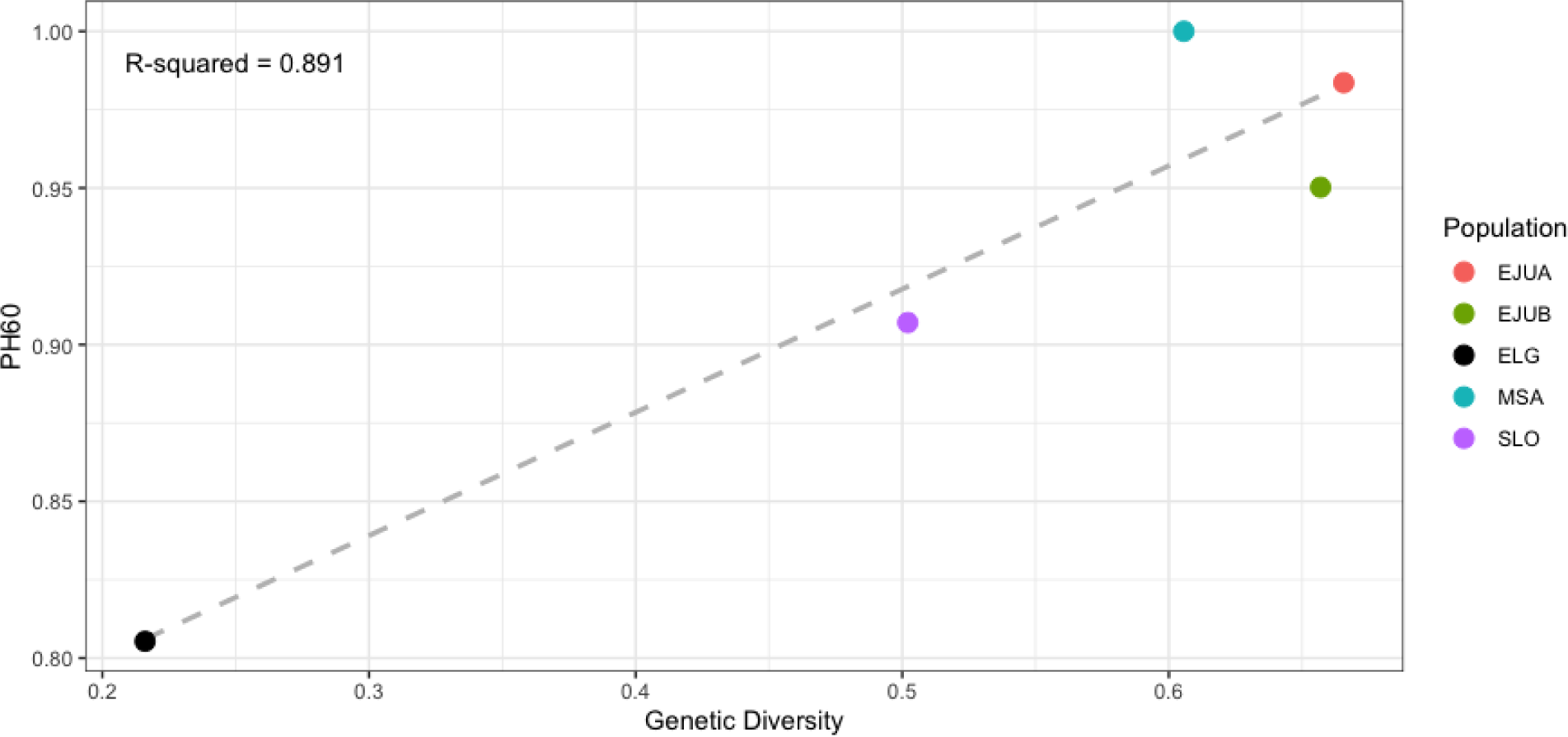
Linear regression between genetic diversity and plant height at 60 days (PH60).

### Genetic Load

The genetic load of the four larger populations (EJUA, EJUB, MSA, and SLO) was calculated using Genomic Evolutionary Rate Profiling (GERP). GERP is a measurement of sequence conservation across multiple species that is based on the number of substitutions at a particular site compared to the neutral expectation. A higher GERP score indicates that there are fewer substitutions than expected at that site, and therefore a derived allele is assumed to be more deleterious. SNP data from 48 individuals (12 from each population) across 7,817 loci were evaluated for deleterious derived alleles. The load was calculated under both an additive and a recessive model, where GERP scores were summed for both heterozygous and homozygous derived alleles under the additive model, and only for homozygous derived alleles under the recessive model. To evaluate the proportion of SNPs that were low, medium, or highly deleterious in each population, loci were grouped into three bins for both models. In the recessive model, the most diverse populations (EJUA and EJUB) had the smallest load compared to the least diverse (MSA and SLO) populations (Figure 4D). These trends were consistent when loci were separated out by degree of deleteriousness (Figure 4A-C). However, these trends were reversed under the additive model (Supplemental Figure 3), where the least diverse populations saw the smallest load compared to those with higher diversity. When only the most deleterious alleles were considered (GERP score > 4), the differences between the populations were not significant. At the intermediate levels (GERP score between 2 and 4), SLO interestingly had the highest load of the four populations, with no significant difference between SLO and EJUA. This suggests that while selection and purging is occurring in all four populations for the most deleterious alleles, SLO lacks the heterozygosity for selection to act upon the intermediately deleterious alleles. The general trends under the additive model, with EJUA and EJUB showing the highest amount of load, suggests masking of deleterious alleles due to heterozygosity at those sites compared to the more homozygous SLO population.

**Figure 4.**
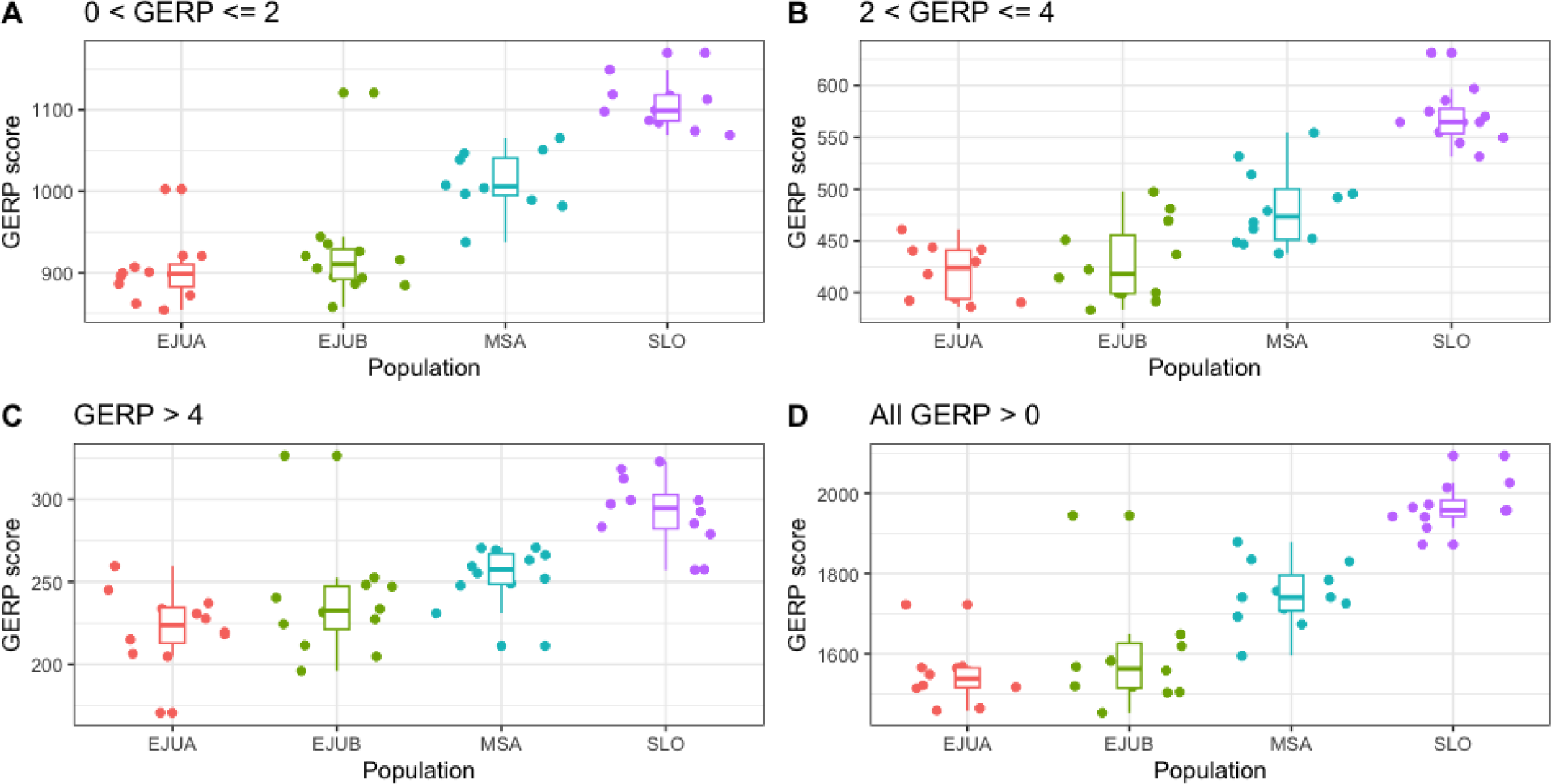
Recessive model for GERP, where only loci with the homozygous derived allele are counted and scores across all loci summed for each individual sampled. A) Lowly deleterious sites B) Moderately deleterious sites C) Highly deleterious sites D) All deleterious sites.

## Discussion

The link between genetic diversity and fitness of a population can be understood in the context of inbreeding depression (Reed, 2003), where a reduction in genetic diversity leads to lower population fitness. Our data suggest that a reduction in genetic diversity at neutral loci can document overall trends of genetic diversity, and in turn, effects of evolutionary or demographic processes on overall fitness. This is observed in the SLO population which appears to have undergone a recent population bottleneck. When comparing fitness with genetic diversity, we saw a positive trend for all traits, which indicates greater genetic diversity within a population results in greater fitness. This was especially apparent in the vegetative traits (PH15, PH30, PH60, NT, and TAB), where the trends of increased fitness correlating with increased diversity were consistent between the populations. The reproductive traits and stomatal conductance showed weaker correlations due largely to ELG having a higher fitness than expected given the low gene diversity of the population. When ELG was removed and the four larger populations were compared, the trends between fitness and genetic diversity became much clearer. ELG, consisting of less than 100 roadside *parviglumis* plants, may be a recent colonization event from the larger EJUA population as ELG alleles were found within the EJUA population. This may explain the instances where ELG had higher levels of fitness in flowering time and stomatal conductance than MSA and SLO despite having lower levels of diversity. Since colonization events only need a few seeds transferred from a nearby population to occur, it is possible that the ELG population saw a dramatic, rapid genetic bottleneck and consequent inbreeding depression. Alternatively, it is also possible that this population managed to become established as it carried alleles for enhanced fitness from EJUA in spite of having lower genetic diversity. This population provides a prime opportunity to study the effects of recent colonization events in future studies.

Structure analysis showed the eastern (MSA, SLO) and western (EJUA, EJUB, ELG) populations grouping together under a k = 2 model, and ELG separated from the other two groups under a k = 3 model. Regional differences appeared between the western and eastern populations in days to flowering and stomatal conductance. These differences may be evidence of divergence between the two populations, as determined by Hufford *et al*. (2010). However, within regions, there was still a positive correlation between fitness and genetic diversity as well as evidence for inbreeding depression in the less fit populations.

With recent reductions in population size, the lower fitness of the SLO and EJUB populations compared to that of MSA and EJUA is cause for concern and does point toward inbreeding depression. The results of our genetic load analysis are also concerning from a conservation standpoint, as the populations with the lowest levels of genetic diversity (and consequently fitness) suffer from the highest levels of genetic load under the recessive model. Levels of load under the additive model indicate the purging of deleterious alleles may not be as effective in the least diverse populations, which again could cause greater inbreeding depression. Populations with higher genetic diversity, and therefore more heterozygosity, can mask deleterious alleles and purge the most deleterious alleles. Additionally, the general trend of lower load in populations with a higher density under the recessive model, and reduced purging of load under the additive model, brings heightened concern for populations experiencing habitat degradation. The reproductive system of *parviglumis* relies on heavy pollen flow traveling a short distance [estimated 39-87m in the EJUA population (Hufford, 2010)] for fertilization. For reproductive success to occur, there must be a high enough population density such that neighboring plants are close enough for pollination to occur. A reduction in population density may lead to lower diversity, lower fitness, and in turn higher genetic load. The potential genetic bottlenecks and resulting inbreeding depression show that we can no longer consider these populations to be stable and that conservation efforts need to be improved and increased. Future ecological and genetic investigations will need to be conducted to assess the state of the *parviglumis* populations in Jalisco.

Both *in situ* and *ex situ* conservation of CWRs should remain a priority to maintain potentially valuable genetic variation for future breeding efforts. With the lack of protected areas and increasing habitat degradation, management of these populations *in situ* will be especially challenging. The genomics era brings many opportunities to explore CWR genomes, especially those in gene banks. However, when it comes to large-scale *in situ* conservation efforts, sequencing and evaluating the genomes of entire populations is not always practical. Our results indicate that there is a much simpler method. It is possible to lead conservation efforts through the use of microsatellite loci by evaluating and managing for heterozygosity. The use of microsatellites with conservation efforts allows for a low-cost and accessible method to manage our remaining CWR diversity and maintain the adaptive potential of these populations.

## Acknowledgements

We thank Michelle Stitzer and Guillaume Ramstein for their suggestions and comments on the GERP analysis, the Iowa State Statistical Consulting group for their help setting up the statistical analyses, and Samantha Snodgrass for providing feedback and encouragement throughout the writing process.

## Scripts and Data Availability

Data for this study and related scripts will be made available upon request and will be deposited into our repository upon publication.

