## Supplemental Figures for "Inbreeding depression leads to reduced fitness in declining populations of wild maize"

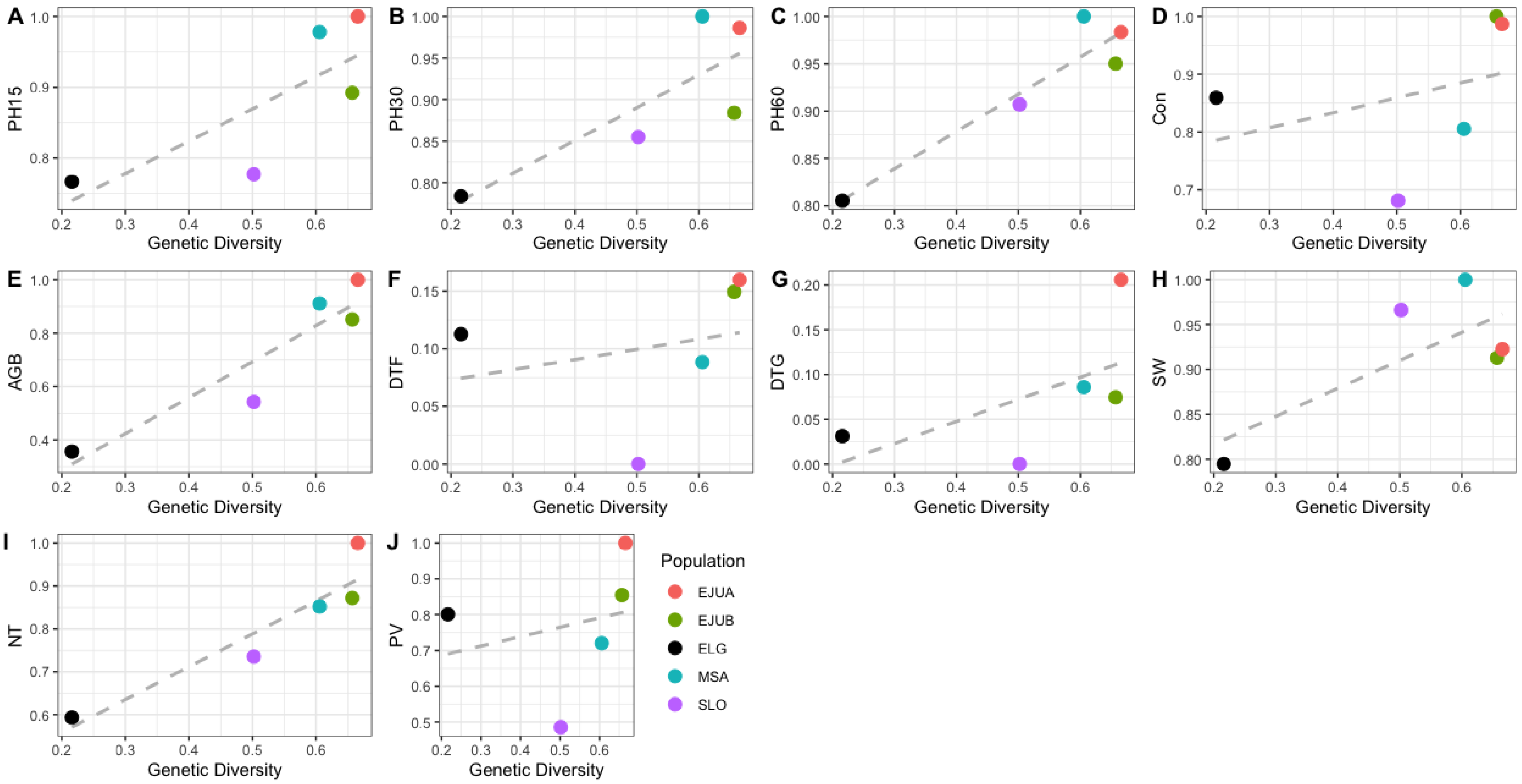


**Supplemental Figure 1.** The relationship between each of the traits measured and genetic diversity, including the ELG population.

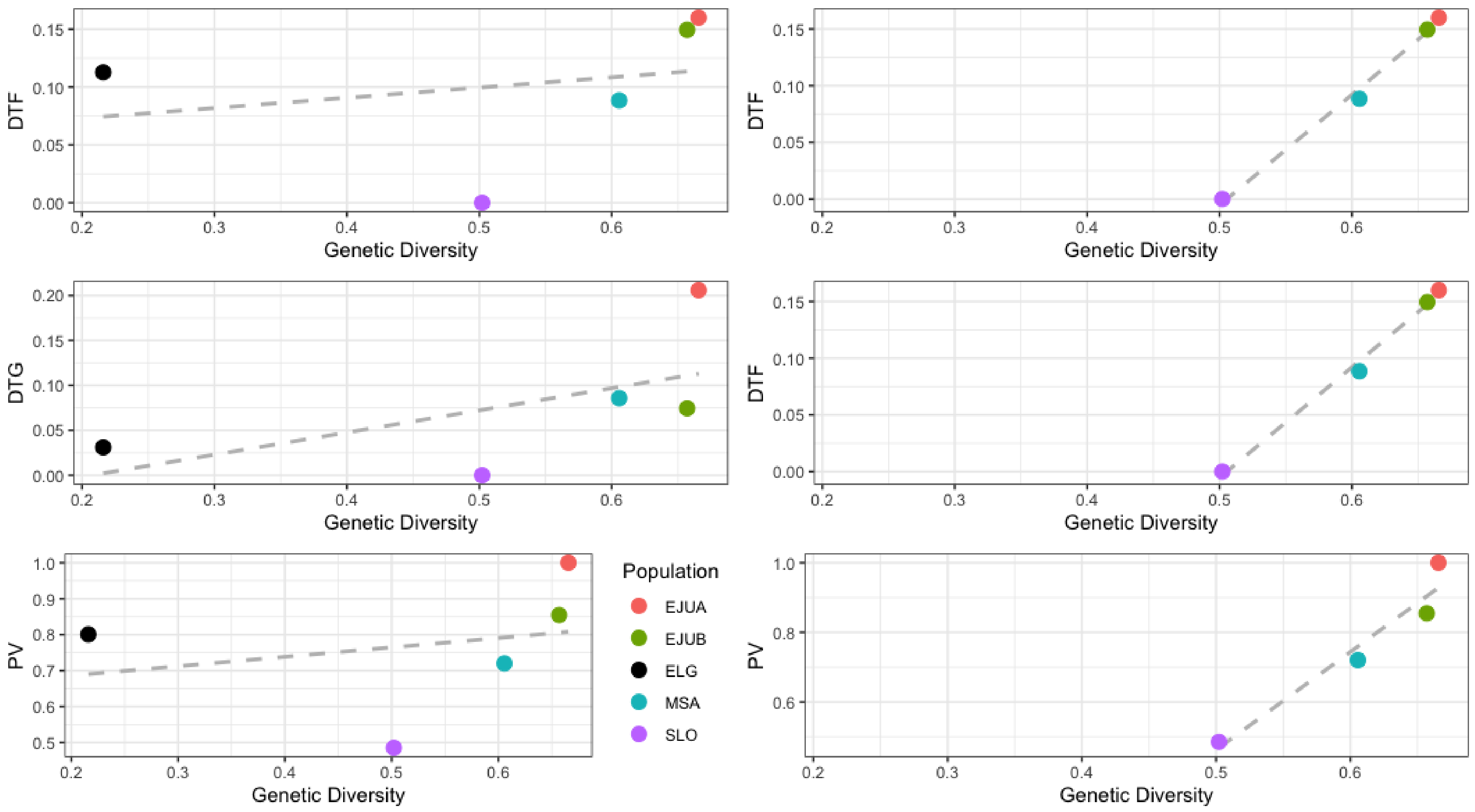


**Supplemental Figure 2.** Excluding the ELG population improved the fit of the linear regression line for DTF, DTG, and PV in relation to genetic diversity.
